# Comprehensive Benchmarking of CITE-seq versus DOGMA-seq Single Cell Multimodal Omics

**DOI:** 10.1101/2021.12.15.472792

**Authors:** Zhongli Xu, Elisa Heidrich-O’Hare, Wei Chen, Richard H. Duerr

## Abstract

The recently developed transcription, epitopes, and chromatin accessibility by sequencing (TEA-seq) and similar DOGMA-seq single-cell trimodal omics assays provide unprecedented opportunities for understanding cell biology, but independent optimization, benchmarking and evaluation are lacking. We explored the utility, pros and cons of DOGMA-seq compared to the bimodal cellular indexing of transcriptomes and epitopes by sequencing (CITE-seq) assay in activated and stimulated human peripheral blood T cells. We identified an optimal incubation time and concentration of digitonin (DIG) for cell permeabilization and found that single-cell trimodal omics measurements after DIG permeabilization were generally better than after an alternative “low-loss lysis” (LLL) permeabilization condition. Next, we found that DOGMA-seq with optimized DIG permeabilization and its ATAC library provides more information, even though its mRNA and cell surface protein antibody-derived tag (ADT) libraries have slightly inferior quality, compared to CITE-seq. Finally, we recognized the additional value of DOGMA-seq for studying lineage-specific T helper cells.

## Introduction

Massively parallel single-cell omic measurements have advanced from single modal measurement of transcriptomes alone; to bimodal, simultaneous transcriptome and cell surface protein epitope measurements (CITE-seq^1^ and REAP-seq^2^), transcriptome and chromatin accessibility measurements (sci-CAR^3^, SNARE-seq^4^, SHARE-seq^5^), and chromatin accessibility and cell surface protein epitope measurements (ASAP-seq^6^ and ICICLE-seq^7^); to trimodal, simultaneous transcriptome, cell surface protein epitope, and chromatin accessibility measurements (DOGMA-seq^6^ and TEA-seq^7^). DOGMA-seq and TEA-seq provide unprecedented opportunities to study complex cellular and molecular processes at single cell resolution, but a comprehensive independent evaluation is needed to compare these new trimodal assays to existing single modal and bimodal assays. In this study, we benchmarked and compared DOGMA-seq under two permeabilization conditions (**Fig. 1A**), as well as DOMGA-seq versus CITE-seq (**Fig. 1B**), using aliquots of the same *ex vivo* tissue culture activated and stimulated human peripheral blood T cell populations.

**Fig. 1.**
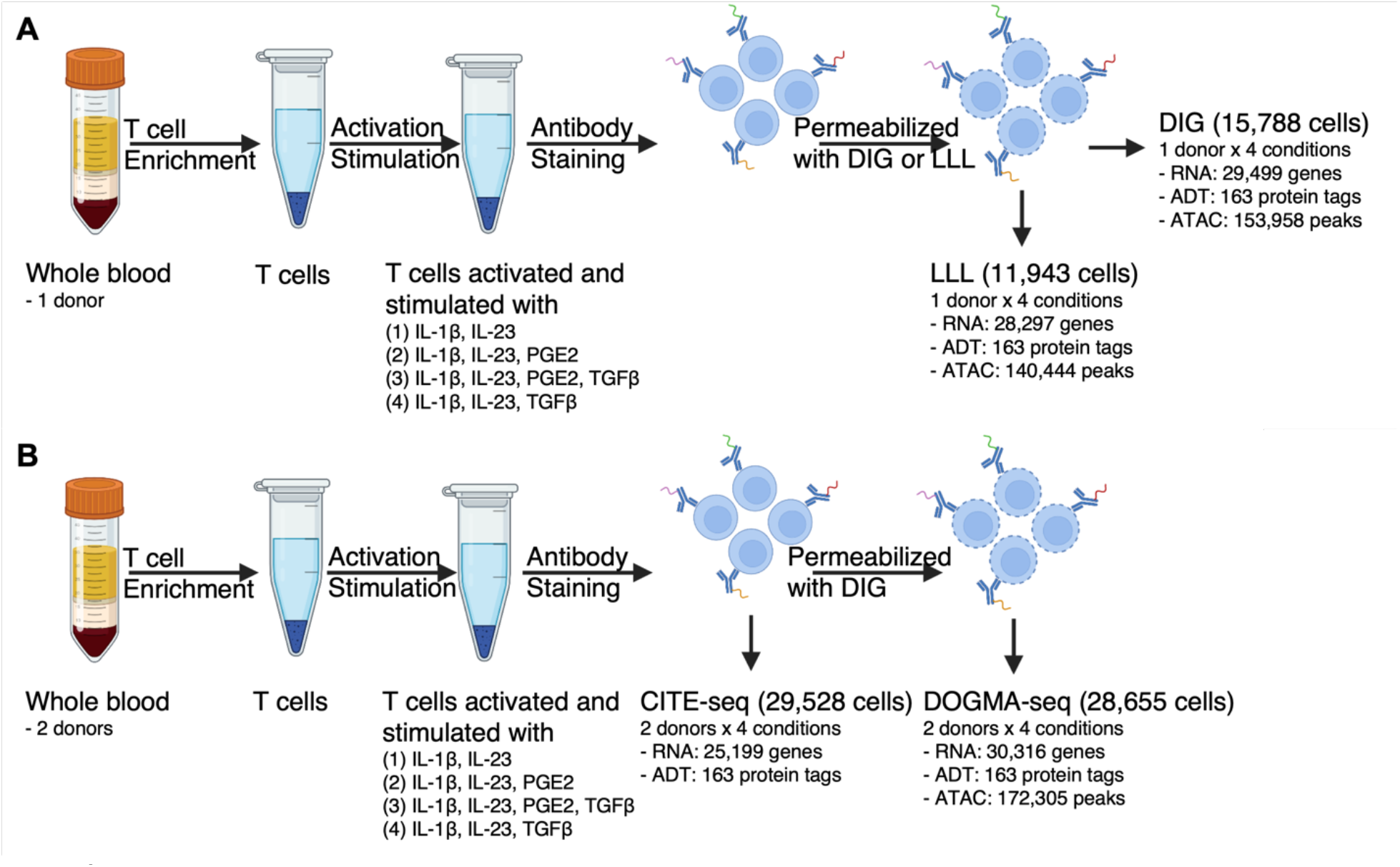
Overview of study designs. (A) Comparison between DIG and LLL conditions. (B) Comparison between CITE-seq and DOGMA-seq.

## Results

The DOGMA-seq and TEA-seq protocols require a cell permeabilization step following hashtag oligonucleotide (HTO) and cell surface protein epitope antibody-derived tag (ADT) labeling to enable preparation of its assay for transposase-accessible chromatin (ATAC) library. Alternative cell permeabilization conditions using either digitonin (DIG) or paraformaldehyde fixation with “low-loss lysis” (LLL) were introduced with the development of the DOGMA-seq^6^ and TEA-seq^7^ assays. Theoretically, DIG permeabilization is milder and will not lyse mitochondrial membranes, because it interacts with cholesterol, and the plasma membrane has a higher cholesterol content compared to the mitochondrial membrane^8^. We optimized the DIG condition used by TEA-seq developers^7^, where cells were incubated with 0.01% DIG on ice for 5 minutes (min), and identified an optimal incubation time and concentration of DIG for permeabilization of peripheral blood mononuclear cells (PBMCs). We had previously determined that incubation with 0.01% DIG for 1 min was sufficient to permeabilize all cells based on acridine orange and propidium iodide (AOPI) staining. We then proceeded to test several dilutions of DIG (0.0025%, 0.005%, 0.0075%, and 0.01%) and found that incubation with 0.0075% DIG for 1 min was an optimal permeabilization condition, since 0.5% of the cells treated with 0.0075% DIG showed red fluorescence indicative of propidium iodide entry into cells with compromised membranes, but lesser concentrations of DIG resulted in lower proportions of permeabilized cells.

Next, we evaluated single-cell trimodal omics measurements after our optimized DIG permeabilization condition compared to the alternative LLL condition (**Fig. 1A)**. We collected PBMC from a healthy donor, enriched untouched T cells, activated and stimulated them under four different *ex vivo* tissue culture conditions, and then split each cell population into DIG and LLL permeabilization groups. Quality control metric comparisons between the two permeabilization groups are shown in the figures (**Fig. 2A-H**). The number of genes detected per cell and TSS enrichment scores are similar after DIG and LLL permeabilization, while DIG resulted in higher protein tag complexity and fraction of mtRNA but a lower fraction of mtDNA. These observations were consistent with those reported by DOGMA-seq developers^6^. However, we observed higher ATAC complexity but a lower fraction of unique molecular identiers (UMIs) mapping to exons after DIG permeabilization, in contrast with the reported observations^6^.

**Fig. 2.**
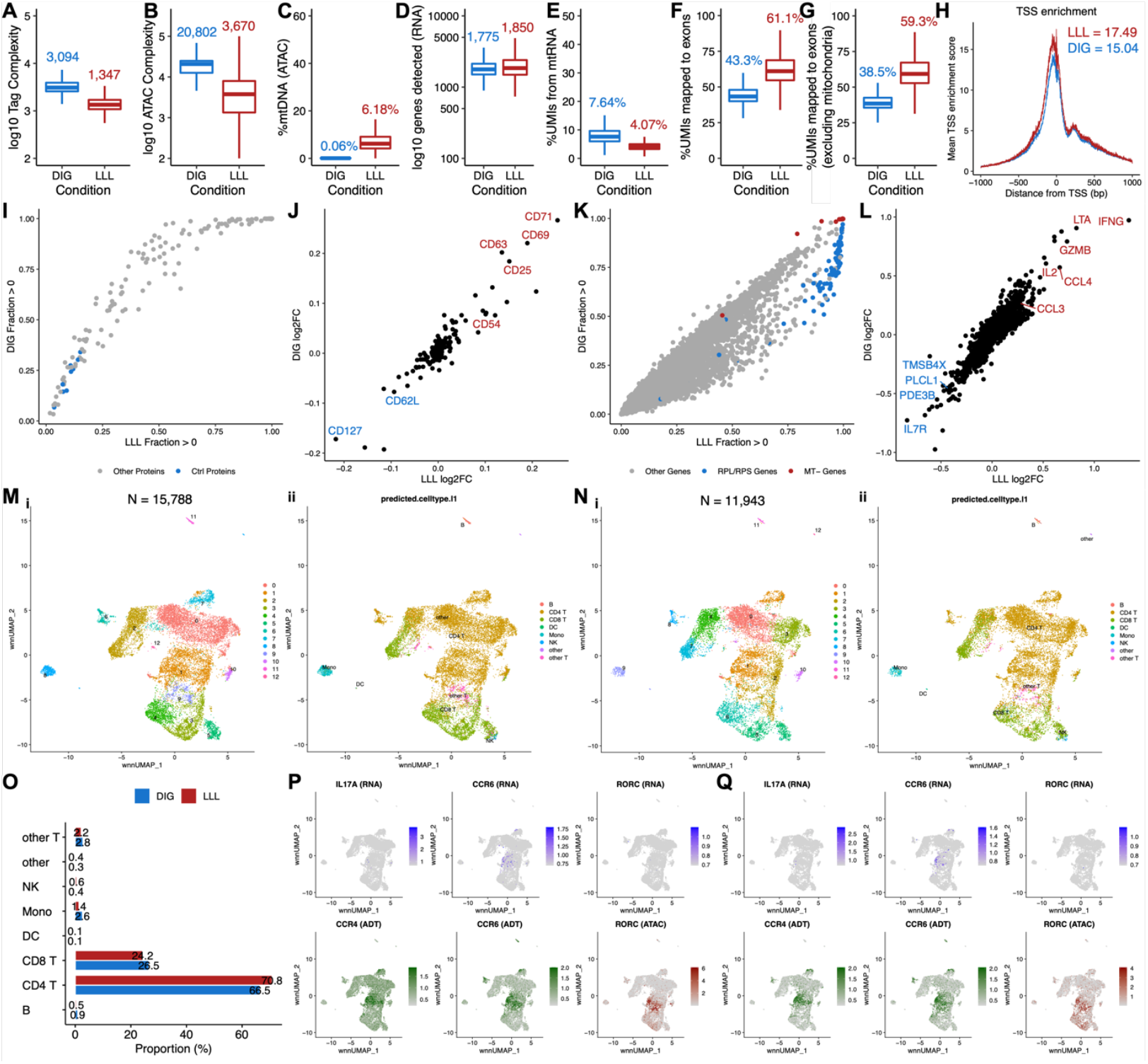
Comparison between DIG and LLL conditions. (A-G) Boxplot showing quality control metric comparisons between DIG and LLL conditions. Median values were indicated with corresponding colors. (A) Protein tag complexity per cell. (B) ATAC fragment complexity per cell. (C) Percentage of ATAC fragments mapped to mtDNA per cell. (D) Number of genes per cell. (E) Percentage of UMIs mapped to mtRNA per cell. (F) Percentage of UMIs mapped to exons per cell. (G) Percentage of UMIs mapped to exons per cell, excluding those mapped to mtRNA. (H) TSS enrichment scores over distance from TSS. Maximum values are indicated with corresponding colors. (I) Pairwise comparison of protein tag detection frequencies under LLL (x-axis) and DIG (y-axis) conditions. Each point represents a single protein tag. Blue points highlight isotype control protein tags. Grey points are all other protein tags. (J) Correlation of protein tag fold change (log_2_) under DIG and LLL conditions in two groups of T cells that were both activated and cultured with IL-1β and IL-23, and one of the two groups was also cultured with PGE2. Top upregulated protein tags are highlighted in red; downregulated protein tags are highlighted in blue. (K) Pairwise comparison of gene detection frequencies under LLL (x-axis) and DIG (y-axis) conditions. Each point represents a single gene. Blue points highlight ribosomal protein genes (RPL/S); red points highlight mitochondrial genes (MT-). Grey points are all other genes. (L) Correlation of gene fold change (log_2_) under DIG and LLL conditions in two groups of T cells that were both activated and cultured with IL-1β and IL-23, and one of the two groups was also cultured with PGE2. Top upregulated genes are highlighted in red; downregulated genes are highlighted in blue. (M)(i) “Harmonized” 3WNN UMAP plot showing clusters identified in 3WNN clustering of DOGMA-seq data under DIG condition. (ii) “Harmonized” 3WNN UMAP plot showing cell types predicted after projection into the Azimuth PBMC reference. (N) (i) “Harmonized” 3WNN UMAP plot showing clusters identified in 3WNN clustering of DOGMA-seq data under LLL condition. (ii) “Harmonized” 3WNN UMAP plot showing cell types predicted after projection into the Azimuth PBMC reference. (O) Bar plot show that proportions of predicted cell types in DOGMA-seq data under DIG and LLL conditions are quite similar. (P-Q) “Harmonized” 3WNN UMAP plots highlight canonical markers for Th17 cells in DOGMA-seq data. ATAC marker is motif activity (the deviations in chromatin accessibility across the set of regions) calculated from ATAC-seq peaks. (P) Under DIG condition. (Q) Under LLL condition.

We calculated the fraction of cells with UMIs > 0 for all 163 protein tags in the TotalSeq-A Universal Cocktail to evaluate protein tag detection rate. As expected, we found higher detection rates for all 163 protein tags after DIG permeabilization (**Fig. 2I**), which is consistent with better preservation of the plasma membrane and cell surface proteins after DIG permeabilization. However, the differences in cell surface protein detection between the two permeabilization conditions didn’t attenuate differential protein tag signals upon differential stimulations at the pseudo-bulk level. For example, when we compared two groups of T cells that were both activated and cultured with IL-1β and IL-23, and one of the two groups was also cultured with prostaglandin E2 (PGE2), we found high correlation of differential protein tag signals between the two permeabilization conditions (**Fig. 2J**).

To evaluate gene detection rates, we calculated the fraction of cells with UMIs > 0 for all 36,601 genes in the reference genome and found slightly higher detection rates for some genes after LLL permeabilization (**Fig. 2K**). By default, the RNA sequencing reads from DOGMA-seq libraries were aligned to both exons and introns by cellranger-arc count. We also aligned RNA sequencing reads from both conditions to exons only or introns only and found that LLL outperformed DIG in gene detection rates when RNA sequencing reads were aligned to exons only (**Extended Data Fig. 1D**), but their performances were similar when the reads were aligned to introns only (**Extended Data Fig. 1E**). To further investigate the difference in gene detection rates, we calculated the proportion of exonic UMIs for each gene and dichotomized genes into exon-dominated genes (genes with proportion of exonic UMIs > 0.5) and intron-dominated genes (genes with proportion of exonic UMIs ≤ 0.5), for DIG and LLL (**Extended Data Fig. 1A and 1B**). We found that genes defined as exon-dominated genes under both conditions had the largest increases in gene detection rates for LLL compared to DIG, whereas genes defined as intron-dominated genes under both conditions had the smallest differences (**Extended Data Fig. 1C**), suggesting that the differences in gene detection rates may be primarily due to differential detection of cytoplasmic genes. We also found that gene fold changes with the addition of PGE2 to the cultures were highly correlated between the two permeabilization conditions at the pseudo-bulk level (**Fig. 2L**).

We next assessed DIG and LLL qualitatively by cluster analysis with the same parameters (including resolution). For both conditions, we performed clustering using RNA, ADT, and ATAC data separately. Slightly more clusters were observed in RNA data under the LLL condition (**Extended Data Fig. 1Gii, Hii**) and in ADT data under the DIG condition (**Extended Data Fig. 1Giii, Hiii**), reflecting their strengths. As expected, more clusters were identified in ATAC data under the DIG condition (**Extended Data Fig. 1Giv, Hiv**), consistent with its higher ATAC complexity, as noted above. We also performed trimodal weighted nearest-neighbor^9^ (3WNN) analysis to leverage all three types of data, and we identified the same number of clusters in DIG and LLL data (**Extended Data Fig. 1Gi, Hi**). To better visualize clusters identified in 3WNN analysis for both conditions, we utilized all three types of data to perform embedding correction using Harmony^10^ and performed automatic cell annotation based on RNA data, using the Azimuth PBMC reference^9^. We found that a few large clusters were not separated in LLL data (**Fig. 2Mi, Ni**), such as the cluster predicted to be enriched for other T cells (**Fig. 2Mii, Nii**). In a head-to-head comparison, the proportions of predicted cell types were quite similar between DIG and LLL (**Fig. 2O, Extended Data Fig. 1F>**). Leveraging ATAC data, we can also locate clusters enriched for Th17 (**Fig. 2P, Q**) and Th1 cells (**Extended Data Fig. 1I, J**) under both conditions, highlighting the value that the ATAC library of DOGMA-seq adds to transcriptome and ADT libraries for the study of lineage-specific T helper cells.

To further investigate the information that DOGMA-seq provides, we performed a head-to-head comparison of DOGMA-seq with our optimized DIG permeabilization condition to CITE-seq (**Fig. 1B**). We collected PBMC from two healthy donors, enriched untouched T cells, activated and stimulated them under four different *ex vivo* tissue culture conditions. After the stimulation experiment, the cells were subjected to CITE-seq and DOGMA-seq, respectively. Using either assay, we observed comparable quality control metrics for protein tag complexity, genes detected, and the fraction of mtRNA (**Fig. 3A-C**). The observations remained unchanged even when the donor difference was considered (**Extended Data Fig. 2A-C**).

**Fig. 3.**
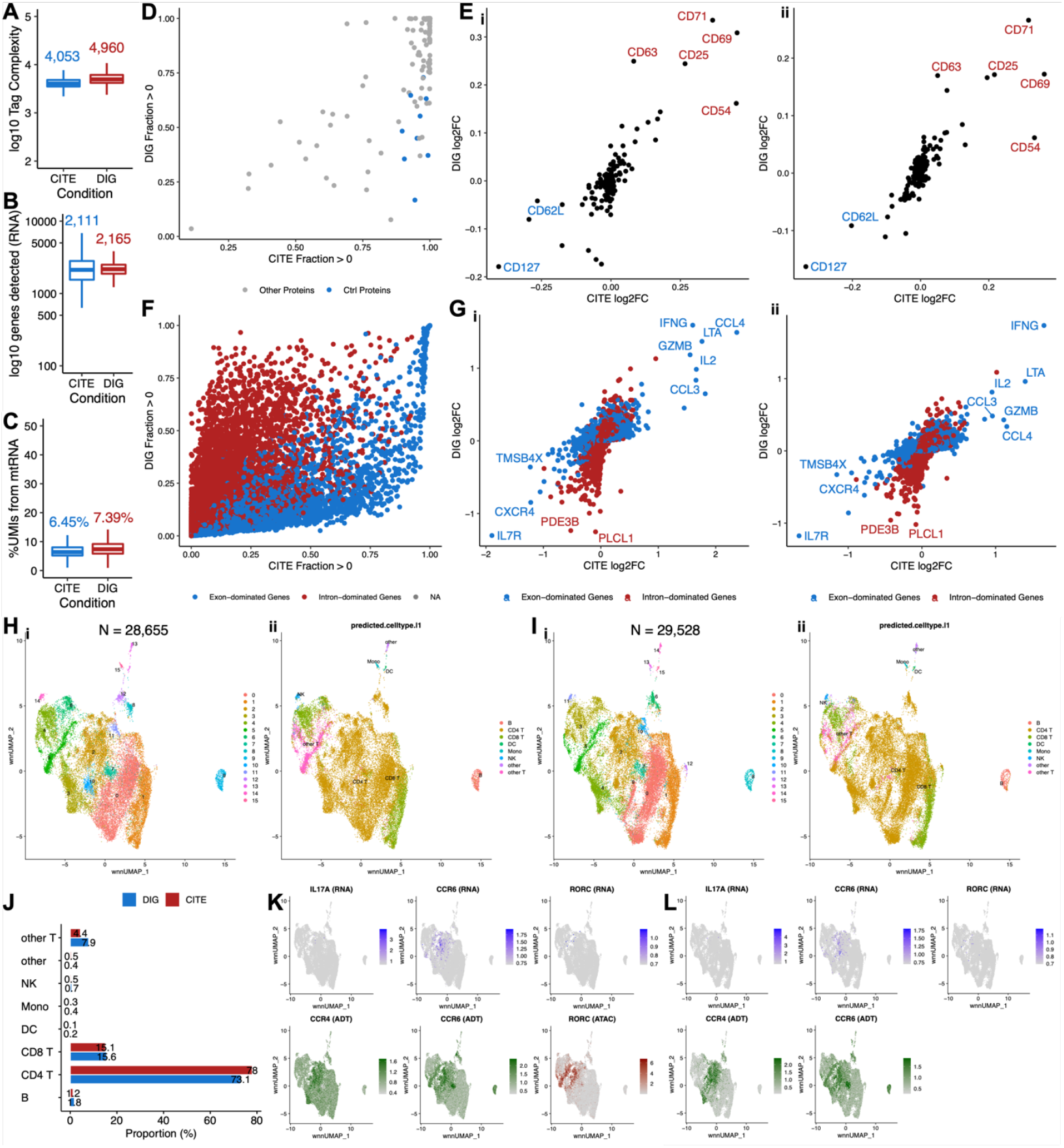
Comparison between CITE-seq and DOGMA-seq. (A-C) Boxplot showing quality control metric comparisons between CITE-seq and DOGMA-seq. Median values are indicated with corresponding colors. (A) Protein tag complexity per cell. (B) Number of genes per cell. (C) Percentage of UMIs mapped to mtRNA per cell. (D) Pairwise comparison of protein tag detection frequencies in CITE-seq (x-axis) and DOGMA-seq (y-axis). Each point represents a single protein tag. Blue points highlight isotype control protein tags. Grey points are all other protein tags. (E)(i) Correlation of protein tag fold change (log_2_) as detected by CITE-seq and DOGMA-seq in two groups of T cells from donor SB775372 that were both activated and cultured with IL-1β and IL-23, and one of the two groups was also cultured with PGE2. Top upregulated protein tags are highlighted in red; downregulated protein tags are highlighted in blue. (ii) in samples from donor SB775393, as (i). (F) Pairwise comparison of gene detection frequencies in CITE-seq (x-axis) and DOGMA-seq (y-axis). Each point represents a single gene. Blue points highlight exon-dominated genes; red points highlight intron-dominated genes. An exon-dominated gene is defined as a gene with proportion of exonic UMIs (DOGMA-seq) > 0.5. An intron-dominated gene is defined as a gene with proportion of exonic UMIs (DOGMA-seq) ≤ 0.5. NA means proportion of exonic UMIs is not available. (G)(i) Correlation of gene fold change (log_2_) as detected by CITE-seq and DOGMA-seq in two groups of T cells from donor SB775372 that were both activated and cultured with IL-1β and IL-23, and one of the two groups was also cultured with PGE2. Selected intron-dominated genes are highlighted in red; selected exon-dominated genes are highlighted in blue. (ii) in samples from donor SB775393, as (i). (H)(i) “Harmonized” 2WNN UMAP plot showing clusters identified in 3WNN clustering of DOGMA-seq data. (ii) “Harmonized” 2WNN UMAP plot showing cell types predicted after projection into the Azimuth PBMC reference. (I)(i) “Harmonized” 2WNN UMAP plot showing clusters identified in 2WNN clustering of CITE-seq data. (ii) “Harmonized” 2WNN UMAP plot showing cell types predicted after projection into the Azimuth PBMC reference. (J) Bar plots show that proportions of predicted cell types in DOGMA-seq and CITE-seq data are quite similar. (K) “Harmonized” 2WNN UMAP plots highlighting canonical markers for Th17 cells in DOGMA-seq data. ATAC marker is motif activity (the deviations in chromatin accessibility across the set of regions) calculated from ATAC-seq peaks. (L) “Harmonized” 2WNN UMAP plots highlight canonical markers for Th17 cells in CITE-seq data.

To evaluate protein tag detection rates, we calculated the fraction of cells with UMIs > 0 for all 163 protein tags in the TotalSeq-A Universal Cocktail and performed a pairwise comparison. As expected, we found that the protein tag detection rates of DOGMA-seq were lower in most cases (**Fig. 3D**), probably due to cell surface protein and/or ADT damage during permeabilization. It is also interesting that the detection rates of isotype control protein tags were higher in CITE-seq. Fortunately, the apparent cell surface protein and/or ADT damage didn’t attenuate differential protein tag signals upon differential stimulations at the pseudo-bulk level. When we compared two groups of T cells that were both activated and cultured with IL-1β and IL-23, and one of the two groups was also cultured with PGE2, significant upregulation and downregulation of a few markers were detectable in both assays and could be replicated in samples from the other donor (**Fig. 3Ei, ii**).

We further compared transcript measurements by CITE-seq and DOGMA-seq. By default, the RNA sequencing reads from the CITE-seq library were aligned to exons only by cellranger multi, whereas those from the DOGMA-seq library were aligned to both exons and introns by cellranger-arc count. This is reasonable since the leakage of transcripts from permeabilized cells can be compensated in some way by additional alignment to introns. We calculated the fraction of cells with UMIs > 0 for all 36,601 genes in the reference genome and found that each assay had its advantage in gene detection rates (**Extended Data Fig. 2G**). We also confirmed that the detection rates of ribosomal genes were lower in DOGMA-seq^7^. We additionally calculated the proportion of exonic UMIs for each gene detected by DOGMA-seq (**Extended Data Fig. 2H**), and dichotomized genes into exon-dominated genes and intron-dominated genes. We observed that CITE-seq had higher detection rates for almost all exon-dominated genes and some intron-dominated genes, whereas DOGMA-seq had higher detection rates for most intron-dominated genes (**Fig. 3F**). Gene fold changes after the addition of PGE2 to the cultures were still correlated at the pseudo-bulk level, but exon-dominated genes and intron-dominated genes formed two lines with different slopes (**Fig. 3Gi**). This finding was replicable in samples from an independent donor using the same protocol (**Fig. 3Gii**). To perform fairer comparisons, we also aligned RNA sequencing reads from both assays to both exons and introns, exons only, and introns only. Generally, CITE-seq outperformed DOGMA-seq in gene detection rates when RNA sequencing reads were aligned to both exons and introns, or exons only (**Extended Data Fig. 2D, E**), but their performances were similar when aligned to introns only (**Extended Data Fig. 2F**).

We next assessed CITE-seq and DOGMA-seq qualitatively by cluster analysis. For DOGMA-seq data, we performed clustering using RNA, ADT, and ATAC data separately (**Extended Data Fig. 3Aii-iv**). We also performed 3WNN analysis to leverage all three types of data (**Extended Data Fig. 3Ai**). For CITE-seq data, we performed clustering for RNA and ADT data separately using the same parameters (including resolution) as those used for DOGMA-seq data (**Extended Data Fig. 3Bii, iii**). Bi-modal WNN (2WNN) analysis was also performed to leverage both RNA and ADT data (**Extended Data Fig. 3Bi**). As expected, fewer clusters and reduced separation were observed in RNA and ADT data of DOGMA-seq compared to CITE-seq since CITE-seq generally had better quality in both RNA and ADT libraries. Despite this, the performance of WNN analysis in DOGMA-seq data was substantially improved by leveraging the additional ATAC library. To better visualize clusters identified in 3WNN analysis of DOGMA-seq data and 2WNN analysis of CITE-seq data, we utilized both RNA and ADT data to perform embedding correction using Harmony^10^. Most clusters identified in 3WNN analysis of DOGMA-seq data could be matched with clusters identified in 2WNN analysis of CITE-seq data, although a few tiny clusters were only separable in either DOGMA-seq or CITE-seq data (**Fig. 3Hi, Ii**). Corroborated by Azimuth PBMC reference projection based on RNA data^9^, the same cell types appeared on matched locations of “Harmonized” 2WNN Uniform Manifold Approximation and Projection (UMAP)^11^ dimensionality reduction plots (**Fig. 3Hii, Iii**) and had comparable proportions in both assays (**Fig. 2J, Extended Data Fig. 2I**), suggesting that Harmony performed well without over-correction. Therefore, we reasoned that biologically meaningful cell types were distinguishable in both 3WNN analysis of DOGMA-seq data and 2WNN analysis of CITE-seq data, as the matchable clusters discussed above. The advantage of DOGMA-seq lies in its simultaneous measurement of chromatin accessibility, enabling the identification of subtypes of T helper cells. Based on expression of canonical markers, cluster 5 in DOGMA-seq data was enriched for Th17 cells (**Fig. 3K**), and cluster 6 was enriched for Th1 cells (**Extended Data Fig. 3C**). These findings were further supported by peaks in genomic regions around canonical markers (**Extended Data Fig. 3Ei-iv**). However, Th17 and Th1 cells were unidentifiable in CITE-seq data (**Fig. 3L, Extended Data Fig. 3D**). Again, these observations highlighted the value of DOGMA-seq to study lineage-specific T helper cells.

## Discussion

In summary, our study optimized the DIG cell permeabilization condition, performed a comprehensive comparison between the DIG and LLL cell permeabilization conditions, and found higher quality ADT and ATAC libraries after DIG permeabilization. We next performed a comprehensive transcriptome and ADT comparison between CITE-seq and DOGMA-seq. We found that DOGMA-seq with optimized DIG permeabilization and its ATAC library provides more information, even though its mRNA and cell surface protein antibody-derived tag (ADT) libraries have slightly inferior quality, compared to CITE-seq. We also recognized the additional value of DOGMA-seq for studying lineage-specific T helper cells. Our study provides a valuable and general guidance for technical and study design considerations for single cell multimodal omics experiments.

## Methods

### Enichment of and Stimulation of Peripheral Blood T Cells

T cells were enriched from 30 ml of whole blood from healthy human subjects, ages 13-35, following the MACSxpress® Whole Blood Pan T Cell Isolation Kit, human (Miltenyi Biotec) protocol. Four aliquots of one million cells each per study subject were stimulated overnight for 12 hours in 1 ml X-VIVO™ 15 Serum-free Hematopoietic Cell Medium (Lonza) with 10 ul ImmunoCult™ Human CD3/CD28/CD2 T Cell Activator (STEMCELL Technologies), 50 ng/ml IL-1B (R&D Systems), and 50 ng/ml IL-23 (R&D Systems), and either 1 uM Prostaglandin E2 (PGE2) (Sigma) or 3 ng/ml TGFB (R&D Systems), or both, for a total of four stimulation conditions per subject. After the overnight incubation, the cells were collected into 15 ml conical tubes through a 30-um strainer and washed with up to 14 ml of PBS/0.2% BSA, centrifuged at 400g for 5 minutes at 4°C, and resuspended in 50 ul Pharmingen Stain Buffer (BSA) (BD Biosciences). An aliquot of each cell suspension was then stained with ViaStain™ AOPI Staining Solution (Nexcelom Bioscience) and viability counts were obtained using a Cellometer Auto 2000 Cell Viability Counter (Nexcelom Bioscience).

### Cell staining with barcoded antibodies

After the addition of 5 ul Human TruStain FcX™ (BioLegend), the cell suspension was incubated for 10 min on ice, and then up to 500,000 cells from each condition were stained with a unique TotalSeq™-A anti-human Hashtag antibody (BioLegend) in 50 ul for 30 minutes at 4°C on a Laminar Wash™ 16-well strip (Curiox Biosystems). The cells were washed in the Laminar Wash Mini System (Curiox Biosystems) with 25 cycles at a flow rate of 10 ul/s. Following collection of each well and a cell count, up to 1 million cells from the different conditions were pooled, centrifuged at 400g for 5 minutes at 4°C and resuspended in 50 ul PBS/0.2% BSA. Fifty ul of TotalSeq™-A Human Universal Cocktail, V1.0 (BioLegend), at a 0.5 dilution, were added to the cell suspension and then the total volume was split between 2 wells of a Laminar Wash™ 16-well strip for a 30 min incubation at 4°C. The cells were washed as previously described, pooled in a 300 ul volume of PBS/0.2% BSA, filtered through a 40 um Scienceware® Flowmi™ Cell Strainer (SP Bel-Art) and counted.

### Single cell preparation for CITE-seq

A total of 30,000 cells were loaded into each of two wells of a Chromium Next GEM Chip G and run on the Chromium Controller (10X Genomics) for Gel Bead-in-emulsion (GEM) generation. The samples were processed according to the CG000204 Chromium Next GEM Single Cell 3’ v3.1 Rev. D protocol (10X Genomics) for RNA library construction while modifications for CITE-seq ADT and cell hashing HTO library preparation were performed according to the TotalSeq™-A Antibodies and Cell Hashing with 10x Single Cell 3’ Reagent Kit v3 or v3.1 single index protocol (BioLegend). The modifications include the addition of ADT and HTO additive primers to a final concentration of 0.2 uM in the cDNA amplification reaction (step 2.2), and using Illumina TruSeq™ Small RNA RPIX (with custom indexes) and Illumina TruSeq™ D70X_short primers, respectively, together with the SI-PCR primer, in ADT and HTO sample index PCRs. The ADT and HTO sample index PCRs were performed on the purified fraction of the supernatant recovered from cDNA amplification cleanup.

### Single cell preparation for DOGMA-seq

The cells were processed according to the CG000338 Chromium Next GEM Multiome ATAC + Gene Expression Rev. D protocol (10X Genomics) for Gene Expression and ATAC library construction while modifications for DOGMA-seq were performed following the “Cell permeabilization with DIG” section of the DOGMA-seq protocol (NYGC Innovation Lab).

We first optimized the concentration of DIG that we would use for the DIG cell permeabilization step. We tested 0.0025%, 0.005%, 0.0075% and 0.01% DIG with about 230,000 cells each, and a DIG concentration of 0.0075% was chosen (see Results).

Approximately 600,000 cells were centrifuged at 400g for 5 minutes at 4°C and permeabilized in 100 ul chilled DIG lysis buffer (20 mM Tris-HCl pH 7.4, 150 mM NaCl, 3 mM MgCl_2_, 0.0075% DIG and 2 U/ul Protector RNase inhibitor (Roche)) for 1 min on ice, followed by the addition of 1 ml of chilled DIG wash buffer (20 mM Tris-HCl pH 7.4, 150 mM NaCl, 3 mM MgCl_2_ and 1 U/ul RNase inhibitor) and pipette mixing, before centrifugation at 500g for 5 minutes at 4°C. The supernatant was discarded, the pellet was resuspended in a very small volume of about 17 ul of diluted Nuclei Buffer (10X Genomics) and 5 ul were used for cell counting.

In a separate experiment, we compared DOGMA-seq’s DIG and LLL protocols. Approximately 600,000 cells were permeabilized with DIG as described above and resuspended in 7.5 ul of Nuclei Buffer. Approximately 300,000 cells were fixed with 0.1% formaldehyde for 5 minutes at room temperature. Glycine was added as a quencher at 0.125M. The cells were washed twice with PBS/1% BSA/RNase inhibitor, centrifuged at 400g for 5 minutes at 4°C and permeabilized in 100 ul chilled LLL lysis buffer (10 mM Tris-HCl pH 7.5, 10 mM NaCl, 3 mM MgCl_2_, 0.1% NP40, 1% BSA, 1 mM DTT and 2 U/ul RNase inhibitor) for 3 min on ice followed by the addition of 1 ml of chilled LLL wash buffer (10 mM Tris-HCl pH 7.5, 10 mM NaCl, 3 mM MgCl_2_, 1% BSA, 1 mM DTT and 1 U/ul RNase inhibitor) and pipette mixing, before centrifugation at 500g for 5 minutes at 4°C. The cells were resuspended in 7.5 ul of Nuclei Buffer. DIG and LLL-treated cells were also processed as described below. The cell recovery after DIG permeabilization was approximately 25-30 %, while the cell recovery after LLL permeabilization was approximately 50%.

Following the CG000338 Chromium Next GEM Multiome ATAC + Gene Expression Rev. D protocol, 30,000 cells were processed for each of two transposition reactions, which were then loaded into each of two wells of a Chromium Next GEM Chip J and run on the Chromium Controller (10X Genomics) for GEM generation.

During sample pre-amplification (step 4.2), ADT and HTO additive primers were added to a final concentration of 0.2 uM. After cleanup and elution in 100 ul, 30 ul were set aside for ADT and 10 ul were set aside for HTO sample index PCR reactions, that were performed using Illumina TruSeq™ Small RNA RPIX (with custom indexes) and Illumina TruSeq™ D70X_short primers, respectively, and the SI-PCR primer.

When generating the RNA library in the Chromium Next GEM Multiome ATAC + Gene Expression protocol, we chose to use the Single Index Plate T Set A instead of the Dual Index Plate TT Set A.

### ADT and HTO primers

ADT additive: CCTTGGCACCCGAGAATT*C*C

HTO additive: GTGACTGGAGTTCAGACGTGTGC*T*C

SI-PCR: AATGATACGGCGACCACCGAGATCTACACTCTTTCCCTACACGACGCTC RPI_SI-GA-A1-1:

CAAGCAGAAGACGGCATACGAGATAGTAAACCGTGACTGGAGTTCCTTGGCACCCGAGAATTC*C*A

RPI_SI-GA-A1-2: CAAGCAGAAGACGGCATACGAGATCCGTTTAGGTGACTGGAGTTCCTTGGCACCCGAGAATTC*C*A

RPI_SI-GA-A1-3: CAAGCAGAAGACGGCATACGAGATGACGCCGAGTGACTGGAGTTCCTTGGCACCCGAGAATTC*C*A

RPI_SI-GA-A1-4: CAAGCAGAAGACGGCATACGAGATTTACGGTTGTGACTGGAGTTCCTTGGCACCCGAGAATTC*C*A

D701_S: CAAGCAGAAGACGGCATACGAGATCGAGTAATGTGACTGGAGTTCAGACGTGT*G*C

D702_S: CAAGCAGAAGACGGCATACGAGATTCTCCGGAGTGACTGGAGTTCAGACGTGT*G*C

D703_S: CAAGCAGAAGACGGCATACGAGATAATGAGCGGTGACTGGAGTTCAGACGTGT*G*C

D704_S: CAAGCAGAAGACGGCATACGAGATGGAATCTCGTGACTGGAGTTCAGACGTGT*G*C

### Library quality control and quantitation

Initial cDNA and ATAC library sizes and estimated concentrations were determined in 2200 TapeStation System High Sensitivity D5000 assays (Agilent Technologies) and initial RNA, ADT and HTO library sizes and estimated concentrations were determined in 2200 TapeStation System High Sensitivity D1000 assays (Agilent Technologies). Qubit™ dsDNA HS assays (Invitrogen), together with average library sizes from the TapeStation assays were used to estimate molarity and to determine library dilutions in the KAPA Library Quantification qPCR assay (Roche Diagnostics), which was used to more accurately estimate molarity.

### Library pooling for sequencing

Libraries from the DOGMA-seq experiment that compared DIG and LLL cell permeabilization were pooled and sequenced with libraries from other projects. The DOGMA-seq DIG and LLL RNA libraries were each represented at approximately 1.99 nM in a pool of libraries that was submitted to the UPMC Genome Center and sequenced using a NovaSeq 6000 S2 Reagent Kit v1.5 (100 cycles) (Illumina) on a NovaSeq 6000 sequencer (Illumina) with a configuration 28/8/0/102 (138 total cycles). DOGMA-seq DIG and LLL ADT libraries were each represented at approximately 2.20 nM and HTO libraries each at approximately 0.27 nM in a pool that was sequenced using a NovaSeq 6000 S1 Reagent Kit v1.5 (100 cycles) with a configuration 28/8/0/15 (51 total cycles). DOGMA-seq DIG and LLL ATAC libraries were each represented at approximately 2.5 nM in a pool that was sequenced using a NovaSeq 6000 S1 Reagent Kit v1.5 (100 cycles) with a configuration 53/8/24/53 and (138 total cycles).

Each of the four other CITE-seq and DOGMA-seq DIG RNA libraries were pooled at approximately 2.5 nM and sequenced using a NovaSeq 6000 S2 Reagent Kit v1.5 (100 cycles) with a configuration 28/8/0/102 (138 total cycles). Each of the four other CITE-seq and DOGMA-seq DIG ADT libraries were pooled at approximately 2.23 nM together with each of the four other HTO libraries at approximately 0.27 nM, and sequenced using a NovaSeq 6000 S1 Reagent Kit v1.5 (100 cycles) with a configuration 28/8/0/15 (51 total cycles). Finally, each of the two other DOGMA-seq ATAC libraries each pooled at approximately 2.5 nM and sequenced using a NovaSeq 6000 S1 Reagent Kit v1.5 (100 cycles) with a configuration 53/8/24/53 (138 cycles).

### Raw sequencing data processing

For CITE-seq, the sequenced RNA, ADT, and HTO libraries were processed and aligned to GRCh38 human reference genome using Cell Ranger software (version 6.1.1) from 10x Genomics, with UMI counts summarized for each barcode. To perform fairer comparisons, we additionally aligned the RNA library to exons only or both exons and introns, without or with the parament --include-introns. To estimate protein tag complexity, ADT library was additionally aligned using the kallisto | bustools workflow (kallisto version 0.46.0, bustools version 0.40.0)^12^. Cell demultiplexing was performed based on HTO libraries using the cellranger multi subcommand. To distinguish cells from the background, cell calling was performed on the full raw UMI count matrices, with the filtered UMI count matrices generated and passed for sample assignment. The outputs assigned per sample were merged for downstream analysis. The size of matrices consisting of singlets was listed below:

RNA: 25,199 genes × 29,528 cells; ADT: 163 protein tags × 29,528 cells; merged from 8 samples (2 donors × 4 conditions)

For DOGMA-seq, the sequenced RNA and ATAC libraries were processed and aligned to GRCh38 human reference genome using an adaption (modified the parameter MAX_CELLS in joint cell calling from 20,000 to 60,000, according to the guidance of 10x Genomics) of the Cell Ranger ARC software (version 2.0.0), with UMI counts, peak counts, and fragments summarized for each barcode. To perform fairer comparisons, we additionally aligned the RNA library to both exons and introns or exons only, without or with the parameter --gex-exclude-introns. To distinguish cells from the background, joint cell calling was performed based on RNA UMIs per barcode and ATAC transposition events in peaks per barcode, with the filtered UMI count and peak count matrices generated. ADT and HTO libraries were aligned using the kallisto | bustools workflow (kallisto version 0.46.0, bustools version 0.40.0)^12^, with the full raw UMI count matrices generated, which were subsequently filtered according to the barcodes in the filtered UMI count and peak count matrices for RNA and ATAC libraries. Cell demultiplexing was performed based on the filtered UMI count matrix for the HTO libraries using the function HTODemux() from the R package Seurat (version 4.0.4)^9^. The size of matrices consisting of singlets is listed below:

DOGMA-seq data under DIG condition:

RNA: 29,499 genes × 15,788 cells; ADT: 163 protein tags × 15,788 cells; ATAC: 153,958 peaks × 15,788 cells; merged from 4 samples (1 donors × 4 conditions)

DOGMA-seq data under LLL condition:

RNA: 28,297 genes × 11,943 cells; ADT: 163 protein tags × 11,943 cells; ATAC: 140,444 peaks × 11,943 cells; merged from 4 samples (1 donors × 4 conditions)

DOGMA-seq data in comparison with CITE-seq:

RNA: 30,316 genes × 28,655 cells; ADT: 163 protein tags × 28,655 cells; ATAC: 172,305 peaks × 28,655 cells; merged from 8 samples (2 donors × 4 conditions)

## Complexity analyses

We adapted the custom code deposited by DOGMA-seq developers (https://github.com/caleblareau/asap_reproducibility/blob/master/global_functions/estimateLibraryComplexity.R) to perform complexity analysis. The code is a translation of the subcommand MarkDuplicates from Picard Tools^13^ and takes the number of unique and duplicate fragments from the Cell Ranger ARC (chromatin) and bustools (protein tag) outputs as inputs. For protein tag, we defined unique and duplicate fragments as described by DOGMA-seq developers^6^. For chromatin, we used the sum of “atac_dup_reads” and “atac_fragments” columns and the “atac_fragments” column from the per_barcode_metrics.csv file.

## Cluster analyses

After the matrices were imported into R, they consisted of singlets that were analyzed using the R package Seurat (version 4.0.4)^9^ and Signac (version 1.3.0)^14^. For the RNA data of either assay, regularized negative binomial regression (SCTransform)^15^ was used to normalize UMI count data, with glmGamPoi^16^ invoked to improve the speed. 3,000 highly variable genes were identified and used in principal component analysis. The first 50 principal components were used in UMAP^11^ dimensionality reduction and clustering using the original Louvain algorithm with resolution = 0.2.

For ADT data of either assay, centered log ratio (CLR) transformation was used to normalized UMI count data. All 163 protein tags were set as variable features and used in principal component analysis. The first 30 principal components were used in UMAP dimensionality reduction and clustering using the original Louvain algorithm with resolution = 0.2.

For ATAC data of DOGMA-seq, Latent Semantic Indexing (LSI)^17^ was used to normalize and reduce the dimensionality of peak count data, with all the peaks selected as variable features. The first 50 (excluding the first component due to correlation with sequencing depth) LSI components were used in UMAP dimensionality reduction and clustering using the SLM algorithm with resolution = 0.2.

For CITE-seq and DOGMA-seq data, we also performed 2WNN (RNA and ADT) and 3WNN (RNA, ADT, and ATAC) dimensionality reduction^9^ using the same numbers of components described above for each modality. Clustering was performed using the SLM algorithm with resolution = 0.2.

To better visualize clusters identified in different assays or under different conditions, we performed embedding correction using Harmony^10^. For the comparison of CITE-seq and DOGMA-seq, we normalized the two data separately as described above and identified variable features for each modality (top 3,000 for RNA and top 100 for ADT). We selected shared variable features between CITE-seq and DOGMA-seq as the variable features used in Harmony. We performed linear dimension reduction and ran Harmony on the linear components for each reduction with assay type as the covariate. UMAP dimensionality reduction was performed using “Harmonized” components for RNA and ADT data, with the “Harmonized” 2WNN UMAP plots generated.

For the comparison of DIG and LLL, we merged the two data sets and normalized them, due to smaller differences between the two data compared with the former comparison. Variable features were identified in the merged data for all three modalities and used in Harmony. We performed linear dimension reduction and ran Harmony on the linear components for each reduction with permeabilization method as the covariate. UMAP dimensionality reduction was performed using “Harmonized” components for RNA, ADT, and ATAC data, with the “Harmonized” 3WNN UMAP plots generated.

We also annotated cells in different assays or under different conditions using reference-mapping approach. We found anchors between our data and Azimuth PBMC reference^9^, with normalization.method = “SCT”, reference.reduction = “spca”, dims = 1:50. Cell type labels were then transferred from the reference to our data.

## Acknowledgements

This research was supported in part by the University of Pittsburgh Center for Research Computing through the resources provided.

## Author contributions

Z.X., E.H., W.C., and R.D. conceived the project and designed the experiments. E.H. performed experiments. Z.X. performed data analysis. Z.X., E.H., W.C., and R.D. wrote the manuscript. W.C. and R.D. supervised the work.

## Data availability

The raw and processed data of CITE-seq and DOGMA-seq, including 8 samples for CITE-seq and 16 samples for DOGMA-seq, will be deposited to Gene Expression Omnibus (GEO) upon acceptance of the paper.

## Code availability

The analysis code will be available in GitHub upon acceptance of the paper.

