## Supplemental Figures for "Comprehensive Benchmarking of CITE-seq versus DOGMA-seq Single Cell Multimodal Omics"

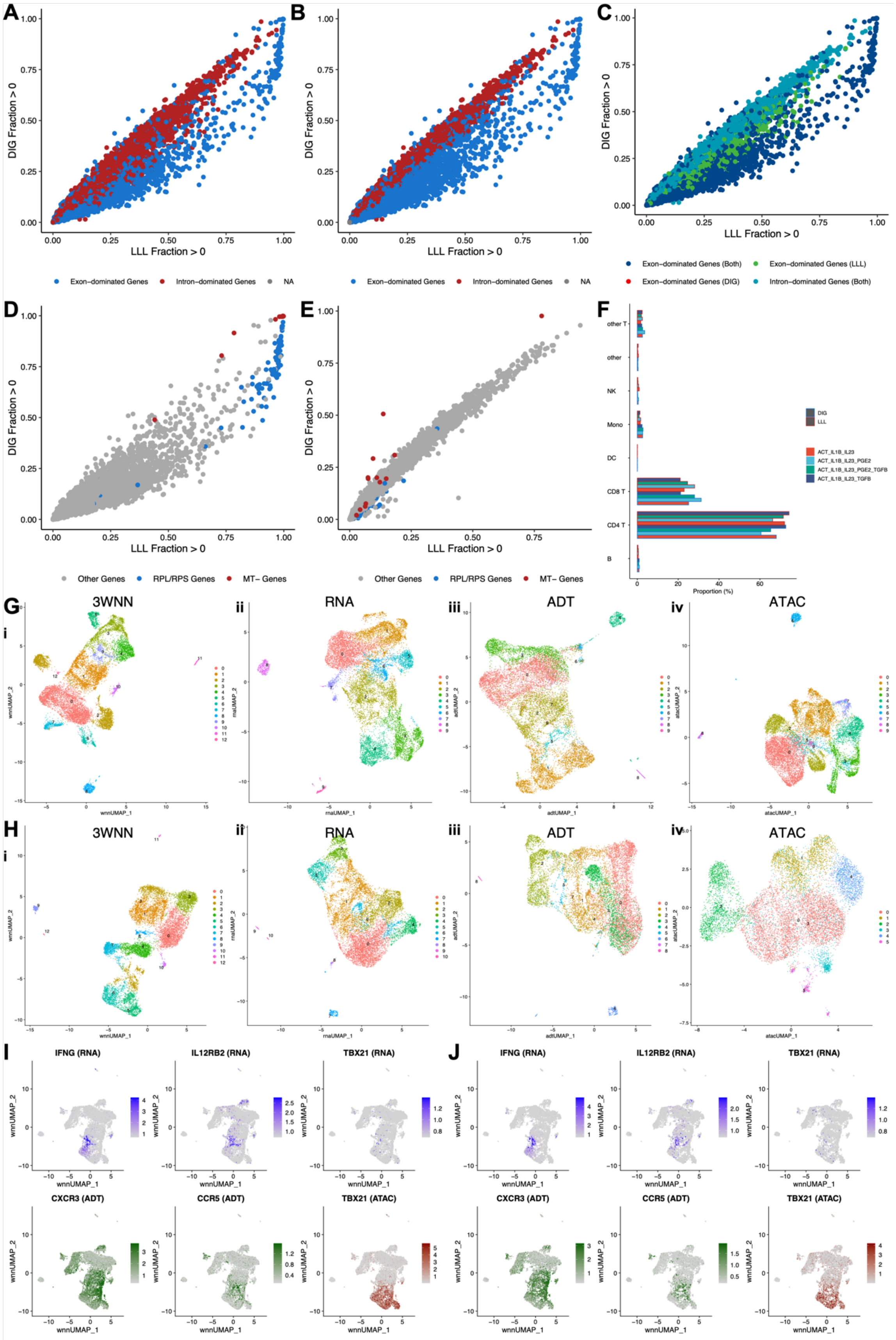

A
B
C
D
E
F
DIG Fraction > 0
LLL Fraction > 0
Exon-dominated Genes
Intron-dominated Genes
NA
Exon-dominated Genes (Both)
Exon-dominated Genes (LLL)
Exon-dominated Genes (DIG)
Intron-dominated Genes (Both)
Other Genes
RPL/RPS Genes
MT- Genes
other T
other
NK
Mono
DC
CD8 T
CD4 T
B
Proportion (%)
DIG
LLL
ACT_IL1B_IL23
ACT_IL1B_IL23_PGE2
ACT_IL1B_IL23_PGE2_TGFB
ACT_IL1B_IL23_TGFB
G
H
i
ii
iii
iv
3WNN
RNA
ADT
ATAC
wnnUMAP_1
wnnUMAP_2
rnaUMAP_1
rnaUMAP_2
adtUMAP_1
adtUMAP_2
atacUMAP_1
atacUMAP_2
I
J
IFNG (RNA)
IL12RB2 (RNA)
TBX21 (RNA)
CXCR3 (ADT)
CCR5 (ADT)
TBX21 (ATAC)

**Extended Data Fig. 1 | Additional comparison between DIG and LLL conditions**

(A-C) Pairwise comparison of gene detection frequencies under LLL (x-axis) and DIG (y-axis) conditions. Each point represents a single gene. Blue points highlight exon-dominated genes; red points highlight intron-dominated genes. NA means proportion of exonic UMIs is not available. (A) An exon-dominated gene is defined as a gene with proportion of exonic UMIs (DIG) > 0.5. An intron-dominated gene is defined as a gene with proportion of exonic UMIs (DIG) ≤ 0.5. (B) An exon-dominated gene is defined as a gene with proportion of exonic UMIs (LLL) > 0.5. An intron-dominated gene is defined as a gene with proportion of exonic UMIs (LLL) ≤ 0.5.

(C) Genes defined as exon-dominated genes under both DIG and LLL conditions are highlighted in blue; genes defined as exon-dominated genes only under DIG condition are highlighted in red; genes defined as exon-dominated genes only under LLL condition are highlighted in green; genes defined as intron-dominated genes under both DIG and LLL conditions are highlighted in cyan.

(D-E) Pairwise comparison of gene detection frequencies under LLL (x-axis) and DIG (y-axis) conditions. Each point represents a single gene. Blue points highlight ribosomal protein genes (RPL/S); red points highlight mitochondrial genes (MT-). Grey points are all other genes. (D) Genes were mapped to exons only under both DIG and LLL conditions. (E) Genes were mapped to introns only under both DIG and LLL conditions.

(F) Bar plot showing proportions of labeled clusters in DOGMA-seq data under DIG and LLL conditions, split by treatment conditions.

(G-H)(i) 3WNN UMAP plot showing clusters identified in 3WNN clustering of DOGMA-seq data. (ii) UMAP plot showing clusters identified based on RNA of DOGMA-seq data. (iii) UMAP plot showing clusters identified based on ADT of DOGMA-seq data. (iv) UMAP showing clusters identified based on ATAC of DOGMA-seq data. (G) Under DIG condition. (H) Under LLL condition.

(I-J) “Harmonized” 3WNN UMAP plots highlighting canonical markers for Th1 cells in DOGMA-seq data. ATAC marker s motif activity (the deviations in chromatin accessibility across the set of regions) calculated from ATAC-seq peaks. (I) Under DIG condition. (J) Under LLL condition.

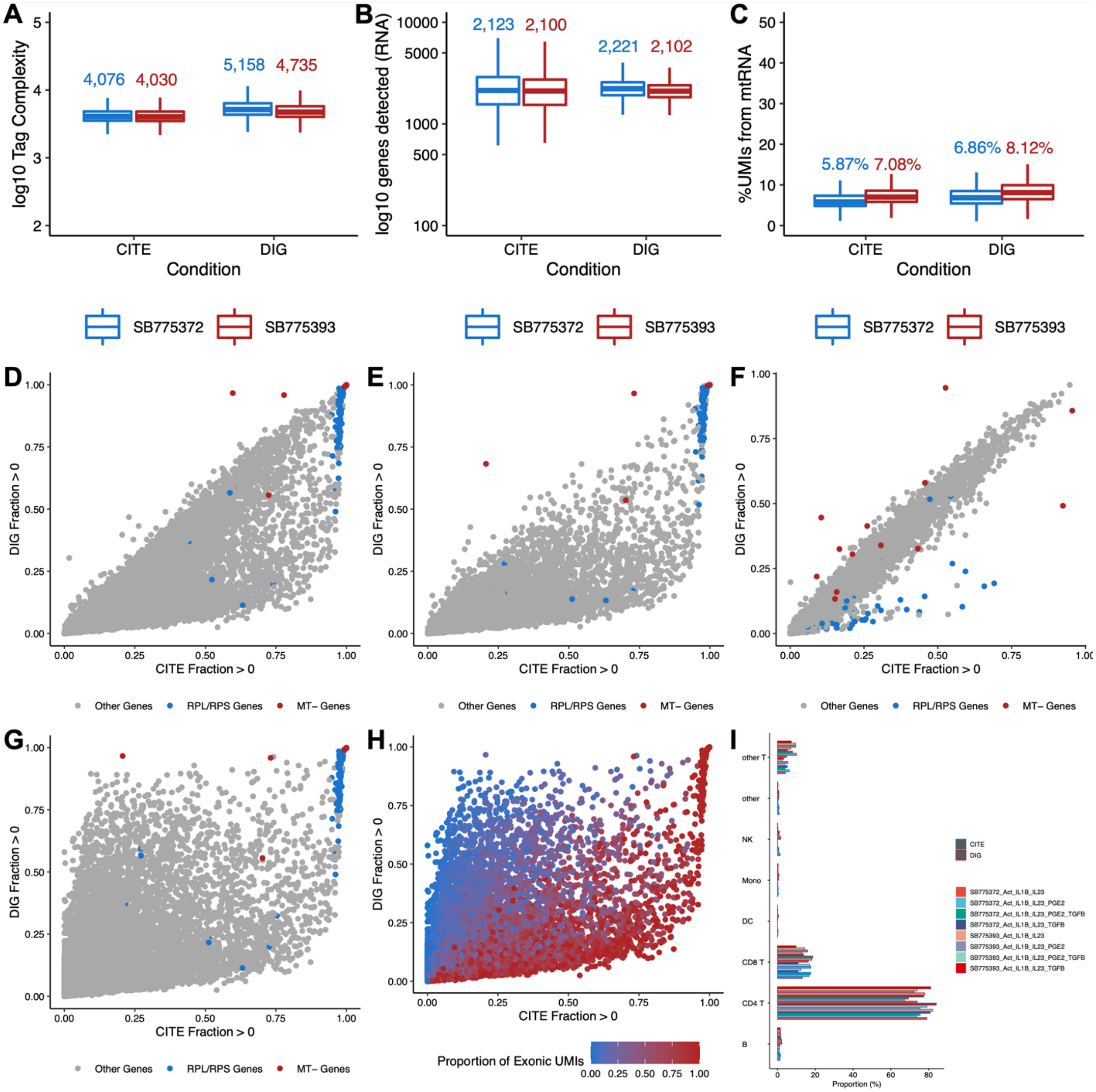


**Extended Data Fig. 2 | Additional comparison between CITE-seq and DOGMA-seq**

(A-C) Boxplot showing quality control metric comparisons between CITE-seq and DOGMA-seq, split by donors (SB775372 or SB775393). Median values were indicated with corresponding colors. (A) Protein tag complexity per cell. (B) Number of genes per cell. (C) Percent of UMIs mapped to mtRNA per cell.

(D-G) Pairwise comparison of gene detection frequencies in CITE-seq (x-axis) and DOGMA-seq (y-axis). Each point represented a single gene. Blue points highlighted ribosomal protein genes (RPL/S); red points highlighted mitochondrial genes (MT-). Grey points are all other genes. (D) Genes were mapped to both exons and introns in both CITE-seq and DOGMA-seq. (E) Genes were mapped to exons only in both CITE-seq and DOGMA-seq. (F) Genes were mapped to introns only in both CITE-seq and DOGMA-seq. (G) Genes were mapped to exons only in CITE-seq, and mapped to both exons and introns in DOGMA-seq.

(H) Pairwise comparison of gene detection frequencies in CITE-seq (x-axis) and DOGMA-seq (y-axis). Each point represented a single gene. The color scale indicated proportion of exonic UMIs (DOGMA-seq).
(I) Bar plot showing proportions of labeled clusters in DOGMA-seq and CITE-seq data, split by donors and treatment conditions.

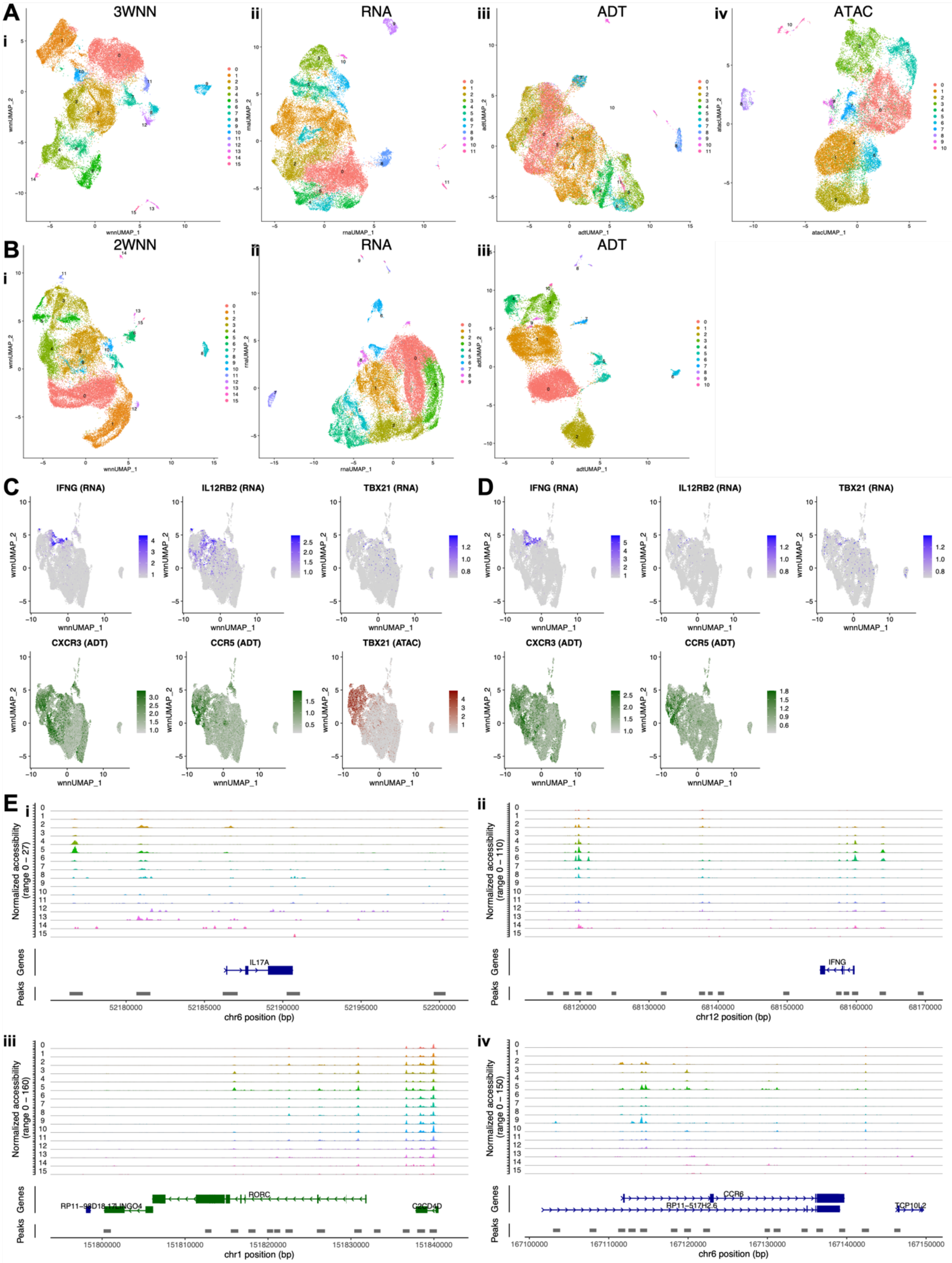


**Extended Data Fig. 3 | Additional comparison between CITE-seq and DOGMA-seq**

(A)(i) 3WNN UMAP plot showing clusters identified in 3WNN clustering of DOGMA-seq data. (ii) UMAP plot based on RNA showing clusters identified in clustering based on RNA of DOGMA-seq data. (iii) UMAP plot based on ADT showing clusters identified in clustering based on ADT of DOGMA-seq data. (iv) UMAP plot based on ATAC showing clusters identified in clustering based on ATAC of DOGMA-seq data.
(B)(i) 2WNN UMAP plot showing clusters identified in 2WNN clustering of CITE-seq data. (ii) UMAP plot based on RNA showing clusters identified in clustering based on RNA of CITE-seq data. (iii) UMAP plot based on ADT showing clusters identified in clustering based on ADT of CITE-seq data.
(C) “Harmonized” 2WNN UMAP plots highlighting canonical markers for Th1 cells in DOGMA-seq data. ATAC marker was motif activity (the deviations in chromatin accessibility across the set of regions) calculated from ATAC-seq peaks.
(D) “Harmonized” 2WNN UMAP plots highlighting canonical markers for Th1 cells in CITE-seq data.
(E)(i-iv) Peaks in genomic regions around *IL17A, IFNG, RORC, CCR6*, canonical markers for Th17 and Th1 cells.
